# Counting of Enzymatically Amplified Affinity Reactions in Hydrogel Particle-Templated Drops

**DOI:** 10.1101/2021.04.21.440664

**Authors:** Yilian Wang, Vishwesh Shah, Angela Lu, Ella Pachler, Brian Cheng, Dino Di Carlo

**Affiliations:** Department of Bioengineering, University of California, Los Angeles, CA; Department of Mechanical and Aerospace Engineering, California NanoSystems Institute, Jonsson Comprehensive Cancer Center, University of California, Los Angeles, CA

## Abstract

Counting of numerous compartmentalized enzymatic reactions underlies quantitative and high sensitivity immunodiagnostic assays. However, digital enzyme-linked immunosorbent assays (ELISA) require specialized instruments which have slowed adoption in research and clinical labs. We present a lab-on-a-particle solution to digital counting of thousands of single enzymatic reactions. Hydrogel particles are used to bind enzymes and template the formation of droplets that compartmentalize reactions with simple pipetting steps. These hydrogel particles can be made at a high throughput, stored, and used during the assay to create ~500,000 compartments within 2 minutes. These particles can also be dried and rehydrated with sample, amplifying the sensitivity of the assay by driving affinity interactions on the hydrogel surface. We demonstrate digital counting of β-galactosidase enzyme at a femtomolar detection limit with a dynamic range of 3 orders of magnitude using standard benchtop equipment and experiment techniques. This approach can faciliate the development of digital ELISAs with reduced need for specialized microfluidic devices, instruments, or imaging systems.

## Introduction

Digital enzyme-linked immunosorbent assays (ELISAs) allow for the measurement of protein biomarkers down to the ultimate sensitivity of single molecules. In contrast to conventional ELISAs performed in ~200 μL reaction volumes, in a digital ELISA workflow, a sample containing a target protein is partitioned into numerous sub-nanoliter microreactors such as microwells or emulsion droplets, such that each partition contains a quantized number of target molecules. If the volume of each partition is small enough, hence the total number of the partitions great enough, many of the microreactions will comprise either 0 or 1 target molecules with a distribution governed by Poisson statistics^1^.

The differentiation of positive partitions (containing 1 or more target molecules) from negative partitions (containing 0 target molecules) requires a signal amplification mechanism associated with the presence of a target molecule inside a partition. Similar to well plate-based ELISA, reporter enzymes such as horseradish peroxidase (HRP), β-galactosidase (β-gal), or alkaline phosphatase (AP) are used to catalytically react with their fluorogenic substrates and linearly amplify signal. Since the signals generated by the reporter enzymes are contained within the original partition, the measurement of target protein concentration is reduced to counting the number of positive partitions containing signals above-background.

While detection of a single β-gal enzyme was achieved as early as 1961 by spraying a mixture of β-gal and its fluorogenic substrate over silicone oil^2^, the reliable measurement of low-abundance protein biomarkers posts two to the partitioning strategy. First, the digital ELISA platform needs to produce and process a large number (~ 10^5^–10^6^)^3,4^ of highly homogenously sized partitions. Monodisperse partition size ensures uniform reaction conditions among all partitions, and thus the signals from the partitions can be directly compared. Second, the partitions need to contain a solid surface to support the immobilized target protein molecules, subsequently formed enzyme-linked immunocomplexes, and sustain repeated washing required for ELISA workflows.

Since microfluidics enabled the fabrication or generation of large numbers of homogeneously sized microscale compartments, digital counting of β-gal enzymes was first demonstrated in microwell arrays^5^ and optical fibers^6^, and matured into full-scale digital ELISA assays in which microwells were loaded with particles^7,8^conjugated with immuno-complexes, leveraging the convenient mixing and washing of micro-particles to expediate the binding of target molecules to their surfaces.^9^ Since the commercialization of microwell-based digital ELISA represented by Quanterix Simoa, new discoveries have been enabled in the diagnosis and mechanistic studies of various diseases^10^ such as for Alzheimer’s disease^11^, cancers^12–14^, cardiovascular diseases^15,16^, cytokine inflammatory markers^17,18^, and tuberculosis^19^. Similar well plate-based platforms include deflectable button microwells^20^ and pre-equilibrium microarrays^21^.

Another design of digital ELISA platforms encapsulates particles into microfluidically-generated and surfactant-stabilized droplets^22,23^. Compared to microwell approaches where the number of microwells are fixed per wafer design and scaling up complicates device handling^24^, the droplet approach scales more freely, since droplets can be produced at up to ~10^6^/s ^22,23^ and as many or few can be utilized as the downstream analysis approach allows. However, using random binding and encapsulation to realize conditions of a single molecule per particle and single particle per droplet, results in <~1% of all droplets containing a single molecule on a single particle, following Poisson statistics. On microwell-based platforms the particle loading efficiency is increased by sizing well cavities for a single particle and infusing an overwhelming supply of particles, however, the majority of them are washed out and uninterrogated^25^.

Another problem shared by microwell and droplet platforms is the cost or operational complexity for end users without microfluidic skills or infrastructure. Both microwells and droplet generators require specialized microfluidic skills to operate or rely on commercial solutions that are specialized and not widely available. Efforts to simplify the partitioning hardware include (1) entirely removing partitioning by depositing signals onto particles *in-situ* with tyramide signal amplification^26,27^ although spacing strategies are required to prevent signal crosstalk amongst particles^28^; and (2) relaxing the monodispersity requirement by statistically compensating for volume polydispersity^29^, adapted to a one-pot reaction^30^.

We present a lab-on-a-particle assay mechanism as a further democratized solution for digital enzyme reaction counting. Hydrogel particles are utilized here both as a solid support to immobilize protein molecules, and as a template for emulsification around the hydrogel matrix. Previously, we and others have shown that spherical particles^31,32^ or particles with tailored surface chemistry^33,34^ can be used to template drop formation to perform measurements of DNA and protein biomarkers. In this work, we performed digital counting of β-gal enzyme on spherical polyethylene glycol (PEG) hydrogel particles templating drops of ~20 pL volume and achieved femtomolar detection limit. The assay workflow is performed solely with standard bench-top equipment and techniques. We envision this lab-on-a-particle mechanism for single enzymatic reaction counting to lay a foundation for democratizing digital ELISA technologies.

## Results

### High throughput production of hydrogel particles

Hydrogel particles used in this work are monodisperse spherical PEG hydrogel particles, composed of 8-arm PEG-vinylsulfone as the main scaffold, crosslinked with DTT with 80% of the thiol crosslinking sites occupied by DTT, and functionalized with biotin-PEG-thiol to introduce biotin moieties and maleimide conjugated fluorophores to impart fluorescence to locate the particles for imaging analysis. Particle production follows our published pH-modulated step emulsification method^35^ (Fig. 1a), yielding droplets with diameters of 25.3 ± 1.4 μm. This step emulsifier is operated at a frequency of 625 KHz, with potential to scale up. The quality of the emulsion remains consistent over a 10-hour production period as shown by the CV of the droplet sizes (Fig. S1).

**Fig. 1.**
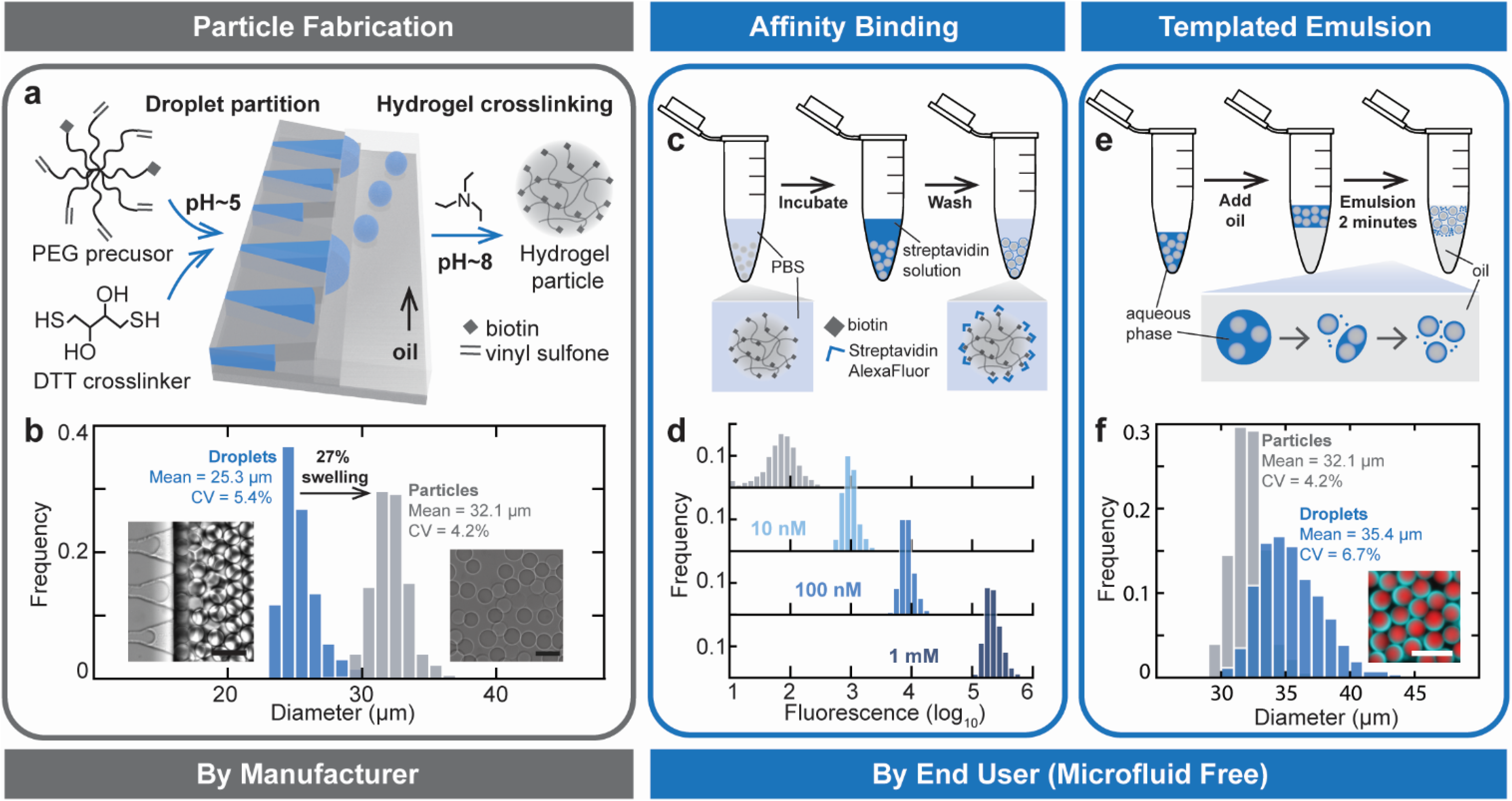
Workflow for enzymatic affinity assays in dropicles. **(a)** Hydrogel particles were produced using step emulsification of polymer precursor droplets at a low non-reactive pH. Droplets containing polymer precursors were crosslinked into hydrogel particles by an increase in pH from 5 to 8. **(b)** Hydrogel particles retrieved from the emulsion droplets (d = 25.3 ±1.4 μm, n = 1,428) swelled 27% in diameter, resulting in hydrogel particles 32.1 ±1.3 μm in diameter (n = 1,122). Bright field microscopy images of droplets and crosslinked hydrogel particles show a change in their refractive index. Scale bar = 50 μm. **(c)** Schematic of affinity binding of fluorescent streptavidin on biotin-modified hydrogel particles following incubation and washing step. **(d)** The fluorescence intensity of particles is correlated to the concentration of the streptavidin solution incubated with particles with a narrow distribution (n = 10,000). **(e)** Schematic of the formation of droplet templated emulsion (i.e. dropicles) created by vigorous pipetting. **(f)** Histograms of size of the dropicles (d = 35.4 ± 2.4 μm, n = 257) compared to the hydrogel particles (d = 32.1 ±1.3 μm, n = 1,122) are shown. Scale bar = 100 μm.

At the droplet collection site, an organic base was introduced in an additional oil phase to modulate the aqueous droplets to pH~8 and induce crosslinking via Michael addition reaction. Crosslinked hydrogel particles were retrieved by breaking the emulsion and washing with hexane and aqueous buffers; after washing the particles go through a 27% swelling in an aqueous suspension and are filtered through a 40 μm cell strainer to removed particles with malformed crosslinks. The resulting hydrogel particles had a mean diameter of 32.1 μm with a 4.2% CV (Fig. 1b).

### Affinity binding on hydrogel particles

Biotin functionalization of particles enables affinity binding to fluorescent streptavidin (Fig. 1c) with a dose response spanning at least 3 orders of magnitude. After incubating 500,000 hydrogel particles with Alexa Fluor 488 tagged streptavidin, bright fluorescent rings were seen surrounding the particles, suggesting streptavidin bound to the biotin moieties on the particle surfaces. (Fig. S2a) 10,000 particles from each incubation were analyzed for the homogeneity of the fluorescent signal, yielding CV values of 7.8%, 5.5% and 3.8% for 10nM, 100nM and 1mM streptavidin concentrations which indicate uniform affinity binding across the particles (Fig. 1d). A linear correlation could be found between the fluorescence of the particles and the concentration of streptavidin, with a nanomolar limit of detection for unamplified direct binding on these particles (Fig. S2b).

### Particle-templated emulsions

Monodisperse droplets templated by hydrogel particles, which we refer to as dropicles, were formed by agitation with an oil and surfactant. When an aqueous suspension of hydrogel particles is mixed with a phase of fluorinated oil and agitated by vigorous pipetting, the shearing from pipetting breaks up the aqueous phase into consequently smaller volumes, until the aqueous layer surrounding a particle is too thin to be further broken up (Fig. 1e, S3). With optimized surfactant constituents the dropicles exhibit a mean diameter of 35.4 μm with a CV of 6.7%, closely approximating the size of the hydrogel particle templates (Fig. 1f). Thus, by simple agitation we were able to create uniform partitions of 20.6±5 pL volume, without the use of microfluidic components such as droplet generators or microwells in forming the compartments.

### Performance of enzymatic affinity assays

We proceeded to perform enzymatic amplification assays on hydrogel particles, following the affinity binding and templated emulsion workflow described above. The particles were first incubated with an enzyme solution to capture the target enzyme molecules, washed, and then emulsified with a solution of a fluorogenic substrate specific to the enzyme (Fig. 2a). When the fraction of the enzyme molecules introduced and bound to particles was below 0.1 of the total particle loading, enzymes were captured on particles in a quantized manner, and digitized fluorescent signals were observed within dropicles.

**Fig. 2.**
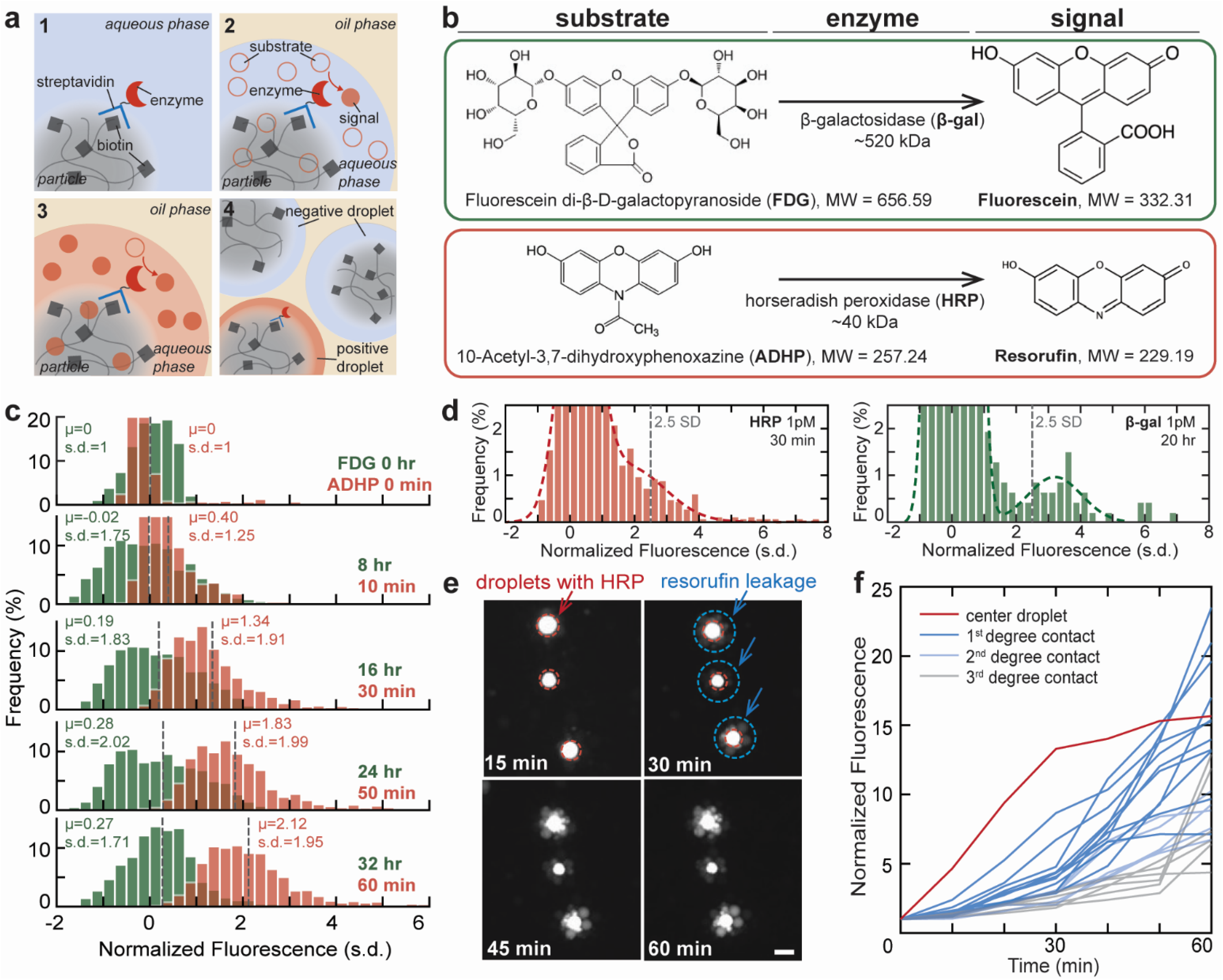
Comparison of enzymatic amplification systems in dropicles. **(a)** Workflow for digital enzymatic amplification assays performed on hydrogel particles. **(b)** Chemical mechanisms for the conversion of fluorogenic precursors to fluorescent molecules for the two enzymatic amplification systems examined on hydrogel particles. **(c)** Comparison of the stability of FDG and ADHP substrates within their respective assay time frame. Signals at later time points were normalized against their measurement at t=0. (n_FDG_ ~3,000, n_ADHP_ ~1,200.) **(d)** Histogram for dropicles after reacting particles with 1 pM of either HRP or β-gal in their respective enzymatic amplification systems after 30 minutes and 24 hours respectively. Signals were normalized against measurement at t=0. A threshold for positive droplets was set to 2.5 standard deviations above the background mean signal. The histograms were fit with a double Gaussian distribution model indicating the resolution of a no signal population and distinct positive population. **(e)** Fluorescence images of resorufin leaking out of the original HRP-containing droplets (marked in red) into nearby droplets (marked in blue) over 60 minutes. Scale bar = 50 μm. **(f)** Fluorescence signals over time for an HRP-containing droplet and nearby droplets over 60 minutes.

We compared the performance of two enzyme systems, evaluating substrate stability, turn-over rate, and signal containment within dropicles. Horseradish peroxidase (HRP) and β-galactosidase (β-gal) are the two most common reporter enzymes used in digital ELISA due to the commercial availability of their fluorogenic substrates (Fig. 2b). HRP catalyzes the conversion of its substrate 10-acetyl-3,7-dihydroxyphenoxazine (ADHP) into resorufin exhibiting bright fluorescence in the 560– 610 nm range. β-gal catalyzes the hydrolysis of β-galactosides through the breaking of the glycosidic bond by a two-step hydrolysis process^36,37^. Multiple fluorogenic substrates of β-gal are commercially available, including those which produce resorufin and fluorescein (e.g. fluorescein-di-beta-galactopyranoside, FDG).

#### Turnover rate

The HRP/ADHP system has significantly higher turnover than the β-gal/FDG system. Resorufin signal generated by HRP can be distinguished within 10 minutes of initiating a reaction in dropicles and plateaus around 30 min (Fig. S4), while sufficient fluorescein signal from the turnover of FDG by β-gal enzyme requires overnight reaction^38,39^.

#### Substrate stability

We measured the chemical stability of FDG and ADHP substrates contained in particle-templated droplets in the absence of enzymes (*i.e.* negative control for enzymatic assays). Due to the difference in turnover rate, droplets containing FDG reaction mix were incubated overnight and imaged at 8-hour intervals, while droplets with ADHP reaction mix were imaged every 10 minutes. The signals from later time points were normalized against their initial readings at t = 0.

While the baseline intensity distribution from the turnover of FDG without enzyme remained similar to their initial readings, spreading slightly in standard deviation (an increase of 0.27 ± 1.71 standard deviations over a 32-hour monitoring window), the intensity distribution from the ADHP reaction mix shifted 2.12 standard deviations higher in intensity on top of a wider (1.95 standard deviations) distribution (Fig. 2c). This can be explained by the photooxidation of ADHP to resorufin^40^ in a peroxidase environment, where the ADHP substrate self-decomposes into resorufin fluorophores unaided by HRP. This baseline level of turnover of the ADHP substrate resulted in a higher noise floor that requires higher levels of enzyme-produced signal to robustly distinguish enzyme containing drops (Fig. 2d).

#### Signal containment

While fluorinated oils ubiquitous in droplet microfluidic assays have very low solubility for large or highly charged biomolecules, dye leakage has been widely reported and attributed to a surfactant mediated transport of small molecules such as fluorophores^41–43^, with resorufin reported to be transported more readily than fluorescein^44,45^. This transport, coupled with the large heterogeneity in enzyme activity within a single population of enzymes, can prevent significant accumulation of fluorescent signals in droplets thereby precluding accurate digital molecular counting.

Since surfactant is necessary for the formation of dropicles, leakage of resorufin fluorophores was significant and could be observed from as early as 15 minutes following compartmentalization and initiation of reactions (Fig. 2e). Fluorescent signals could be detected at even higher levels from satellite droplets surrounding a positive particle as more time passed, up to 60 minutes. Further analysis shows while the reaction in the center HRP-containing droplet plateaus after 30 minutes, signals from the neighboring droplets continue to increase, likely as a combined consequence of dye leakage and the baseline turnover of resorufin molecules.

To decouple enzymatic activity from fluorophore leakage, we observed the transport of fluorescein and resorufin dyes loaded in dropicles by emulsifying particles with fluorescein and resorufin solutions and tracking the signal changes over time. Normalized based on photobleaching, the loss of resorufin fluorophores was rapid with >80% loss of signal within the first 30 minutes of reaction, while the leakage of fluorescein reached e quilibrium after 20 minutes with the majority of the fluorophores remaining within the dropicle (Fig. S5).

Weighing turnover rate, substrate stability, and signal containment, although the HRP/ADHP system had a higher turnover rate leading to a shorter incubation time, the high baseline turnover of ADHP substrate without enzyme and the transport of converted resorufin signaling molecules out of dropicles are both expected to reduce the accuracy of measurements. We therefore decided to use the β-gal/FDG enzyme system for single enzymatic reaction counting on hydrogel particles with an overnight incubation for the enzymatic amplification step.

### Enzymatic affinity assay for digital molecule counting

We performed digital counting of β-galactosidase enzyme molecules. Hydrogel particles were incubated with streptavidin labeled β-gal at concentrations from 2.5 fM to 4 pM, washed, and then emulsified with FDG substrate mixture. After a 20-hour incubation, dropicles were transferred to a reservoir for imaging and analysis. Droplets exhibiting fluorescent signals >2.5 standard deviations higher than the mean intensity of negative control droplets were counted as positive (i.e. containing 1 or more molecules of β-gal, Fig. 3c). Around 50,000 dropicles were imaged from each reaction, and 3 repeats were performed for each β-gal concentration.

**Fig. 3.**
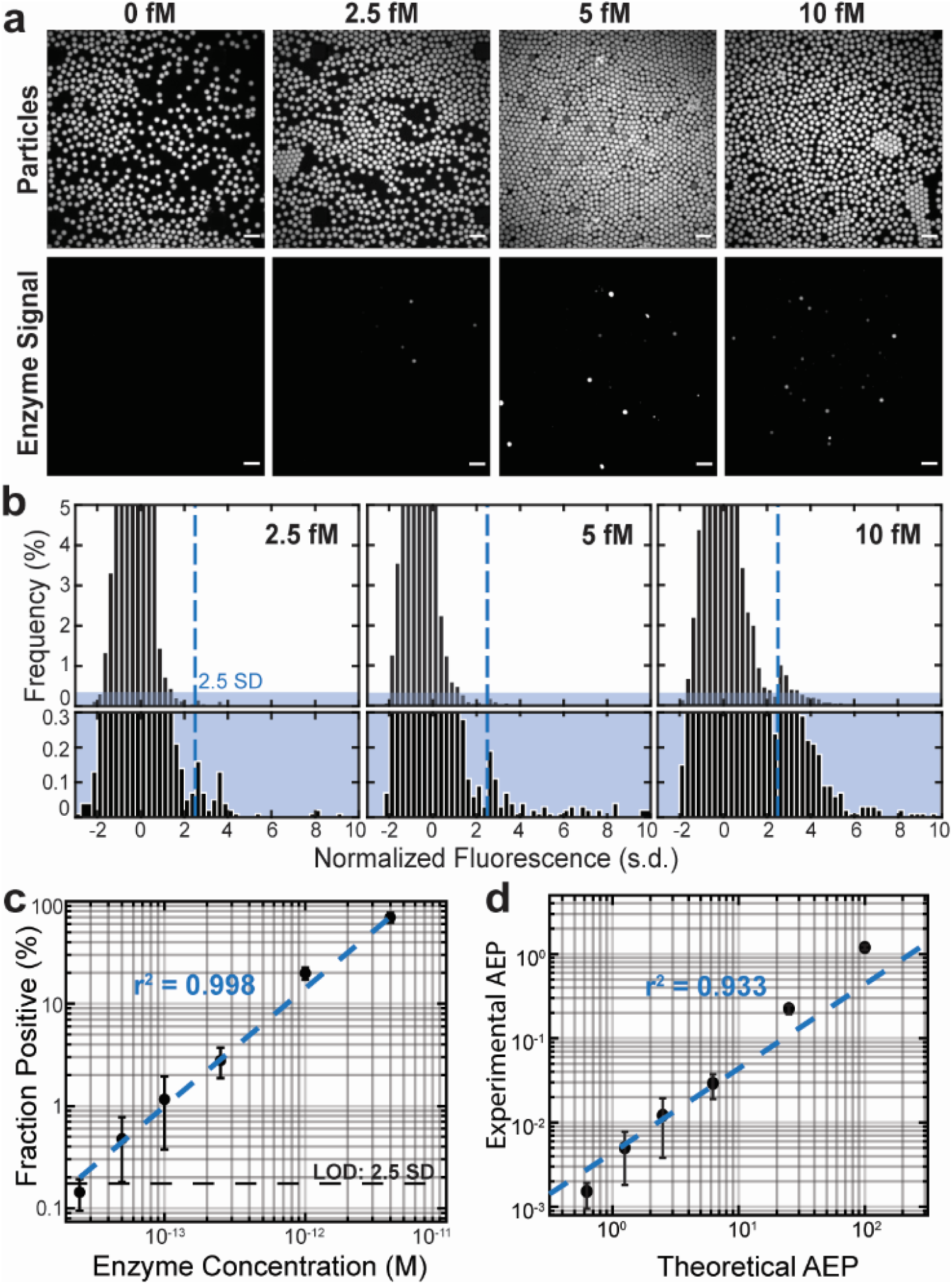
Digital counting of β-gal enzymes in dropicles. **(a)** Fluorescence microscopy images of particles (TRITC) incubated with β-gal enzyme at difference concentrations, and the digitalized fluorescein signals (FITC) from the dropicles indicating events of enzymatic amplification. Scale bar = 100 μm. **(b)** The fraction of positive dropicles (defined by FITC signals >2.5 SDs above the mean background fraction) correlates linearly to the concentration of β-gal enzyme loaded onto particles (r2 = 0.998, n ~ 50,000, N = 3). We observe a 3-order of magnitude dynamic range with a limit of detection at 2.5 fM. **(c)** The experimental average enzyme per particle (AEP) correlates linearly with theoretical values across the range of the experimented β-gal concentrations (r2 = 0.933, N = 3). **(d)** Distribution of fluorescence signals in dropicles from particles incubated with 2.5 fM, 5 fM and 10 fM β-gal solutions. Positive droplets (defined by signals >2.5 SDs above the mean background intensity) increased in frequency with higher β-gal concentrations.

The fraction of positive droplets showed a linear correlation with the starting β-gal concentration over 3 orders of magnitudes (r^2^ = 0.998). The limit of detection for β-gal enzyme counting was 2.5 fM (Fig. 3a). By back calculating from the fraction of positive droplets per experiment to average number of enzymes per particle (AEP) following Poisson statistics, we observed a linear correlation between the experimental average enzyme per particle (AEP) and the theoretical AEP with r^2^ = 0.933 (Fig. 3b).

### Enhancing binding using active absorption of sample

We take advantage of the absorption of fluid by the hydrogel particles once dried to facilitate binding of enzyme driven to the particle surface by the rehydrating flow (Fig. 4a–b). Due to the high weight fraction (~90%) of water in the volume of our hydrogel particles they shrink in size significantly when undergoing a drying process and swell when rehydrated, drawing fluid into their interior. The diameter of hydrogel particles increased from 16.3 ± 1.6 μm to 30.3 ± 2.4 μm during rehydration (Fig. S6). This rehydration process completes within ~2 seconds for each particle (Video S1), resulting in a 6.4x volume increase (Fig. 4c). This corresponds to a calculated convective flow to the surface of ~7 micrometers/sec.

**Fig. 4.**
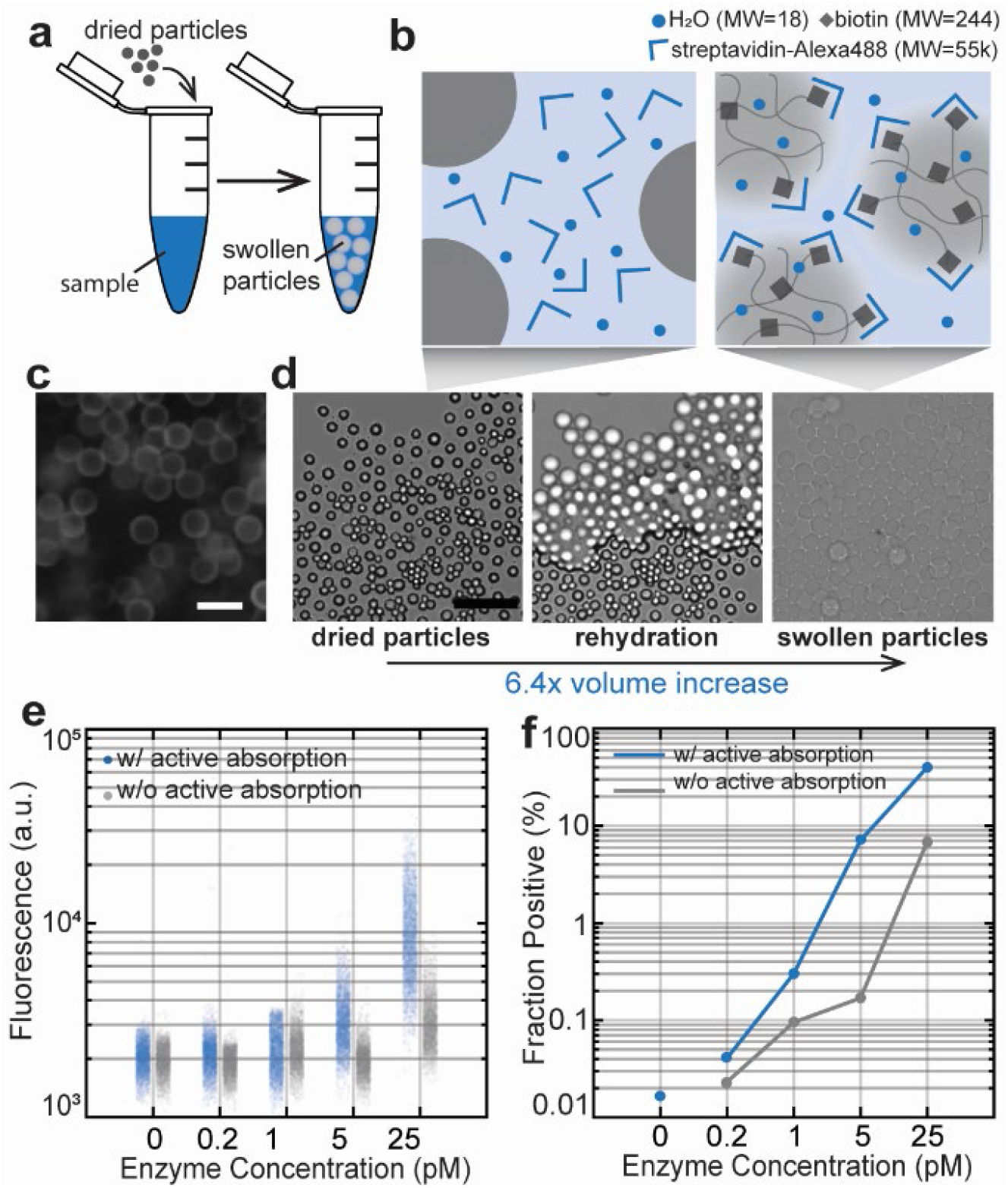
Enzymatic amplification assays facilitated by active absorption by hydrogel particles. **(a)** Schematic of the experimental operations for active absorption using dried hydrogel particles. **(b)** Schematic of the mechanism of active absorption during rehydration of hydrogel particles with sample solution. **(c)** Fluorescence microscopy images showing streptavidin-Alexa Fluor 647 binding to hydrogel particle surfaces following active absorption. Bright fluorescent rings around the particles shows streptavidin did not penetrate the hydrogel particle matrix due to the hydrogel’s limited porosity. **(d)** Bright field microscopy images showing dried hydrogel particles swelling in volume during rehydration, resulting in a 6.4x volume increase. Scale bar = 50 μm. **(e)** The fluorescence signal at a given concentration of enzyme and **(f)** the percent positive dropicles increased with active absorption (n ~ 50,000).

Swelling particles absorb water and other aqueous content into their hydrogel matrix, but the porosity of the hydrogel material prevents larger molecules and proteins from penetrating the matrix. This can be visualized on the hydrogel particles used in this work by the rapid formation of fluorescent rings when particles are incubated with fluorescent streptavidin (MW = 55kDa) (Fig. 4d). This behavior contrasts with the homogenous fluorescence across the particles when filling with small dye molecules (MW = 244Da) not attached to proteins. As a result, during the rehydration process when the hydrogel particles absorb water and swell in size, larger molecules such as streptavidin and enzymes are driven onto their surface, achieving higher local concentrations and effectively facilitating binding to particles (Fig. 4b).

We performed a side-by-side comparison of unfacilitated binding with binding facilitated by active absorption for digital counting of β-gal. Various concentrations of β-gal dilution were added to the same numbers of hydrated and dried hydrogel particles and incubated on a rotating rack for the same amount of time. The particles were then prepared for the standard FDG/fluorescein compartmentalization, signal generation, and imaging. We observe increased sensitivity for assays facilitated with active absorption binding, indicated by an increase in the fraction of positive droplets for the same concentration (Fig. 4e). For concentrations beyond the digital counting regime, the increased binding yielded an increase in the fluorescence signals among all droplets in the same reaction. This indicates a potential to leverage the volume elasticity of hydrogel particles to effectively increase sample concentration, and lower limits of detection.

## Discussion

### The hydrogel particle-based workflow

In this work we demonstrated a lab-on-a-particle approach to digital counting of single enzyme molecules, leveraging the hydrophilicity of hydrogel particles to act as wick to drive flow to the particle surface, the solid particle surface as a support for immunocomplexes and a boundary for uniform droplet generation with random mixing. The combination of these features effectively reduces the need for microfluidic components for the creation of homogeneously sized partitions. Thus, the burden of microfluidics falls solely on the initial generator of hydrogel particles. Once supplied particles, the end user can perform digital affinity assays with standard benchtop techniques like pipetting and centrifugation.

Besides a streamlined workflow, the hydrogel particle approach also overcomes the limit of double Poisson loading, since the process of templated emulsion is not governed by Poisson statistics. In fact, leaving as little aqueous solution as possible to the particle suspension before emulsification is preferred for better emulsion results. We have also found the reliability of the emulsion results depends on the intensity and length of agitation, the volume ratio of the aqueous and oil phases, and the surfactant constituents in aqueous and oil phases. Additionally, this approach to batch emulsification and processing is highly scalable, where to increase the number of desired droplets, the user only needs to increase the number of template particles.

The hydrogel materials of these particles allow the physical and chemical properties of the particles to be customized for the specific need of the assays. The biotin conjugation moieties in this work could be swapped into other common functional groups such as succinimides, aptamers, nucleic acids, and antibodies. Multiplex labeling and barcoding could be integrated into the particle matrix. The porosity and rigidity of the particles could also be controlled by the selection of hydrogel matrix.

### Digital enzyme counting on particles

When demonstrating digital counting of β-gal enzymes with enzymatic signal amplification in particle templated emulsions, we noticed a reduced experimental compared to theoretical AEP. Similar observations were reported in other studies on digital protein counting^7,26,46^. Since a linear correlation was observed between the experimental and theoretical AEPs regardless of the shift, we hypothesize that this shift reflects either a loss of molecules when streptavidin-β-gal was incubated with particles for binding, or amid the agitation process during emulsification. The increase in sensitivity achieved by using active absorption supports this interpretation that loss of target molecules occurs during the incubation step.

Besides loss of target molecules, another factor limiting the sensitivity of particle-based enzyme counting is the capacity of imaging analysis. Although 500,000 dropicles were formed, only ~50,000 ended up imaged with the current microscopic set up. Limits of detection and quantitative accuracy could be improved if the emulsified particles are analyzed with high-throughput flow cytometers capable of processing 500,000 or more dropicles, as this will help decrease the Poisson noise from false positive counting^8,9,47^. These cytometers are often equipped with more sensitive detectors like photomultiplier tubes that will also help further distinguish positive and negative signals. While others have analyzed double emulsions in flow cytometers^48,49^ since dropicles are supported by a hydrogel particle core they may be more likely to sustain fluid shear stresses in a flow cytometer that can operate with an oil continuous phase, for example the On-Chip Sort^50^. Ultimately, we envision the hydrogel particles can be produced at centralized sites and conveniently distributed to enable highly sensitive immunoassays for various biomarkers. No significant changes in the physical and chemical properties of the particles were observed when stored for up to 3 months at 4°C in a buffered aqueous solution. We have also explored alternative storage conditions such as ethanol and lyophilized. Together these capabilities lay the groundwork for a minimally-instrumented and accessible “lab-on-a-particle” solution to the analysis of molecules^51^ and cells^52^ at the ultimate limits of biology.

## Experimental

### Device fabrication

#### Step emulsifiers

Step emulsifier microfluidic droplet generators were fabricated using soft lithography. Master molds were fabricated on mechanical grade silicon wafers by a two-layer photolithography process with KMPR 1010 and 1050 (MicroChem Corp), the first and second layers defining the height of the nozzle channel and the reservoir region respectively. The nozzle channel measured 700 μm (L) × 20 μm (W) × 7.2 μm (H), and the droplet collection reservoir measured 4 cm (L) × 2 mm (W) × 80 μm (H) and were confirmed using a Veeco Dektak 150 Surface Profiler. Devices were molded from the masters using PDMS Sylgard 184 kit (Dow Corning). The base and crosslinker were mixed at a 10:1 mass ratio, poured over the mold, degassed, and cured at 65 °C overnight. PDMS devices and cleaned glass microscope slides (VWR) were then activated via air plasma (Plasma Cleaner, Harrick Plasma) to form permanent bonding. The bonded devices were then treated with Aquapel for 1 min and rinsed with Novec 7500 oil (3M) to render the channels fluorophilic. After surface treatment, devices were placed in an oven at 65 °C for 1 h to evaporate the residual oil in the channels.

#### Imaging reservoirs

To image dropicles and particles, imaging reservoirs sizing 5 cm (L) × 3 cm (W) × 50 μm (H) were fabricated using the same soft lithography techniques, allowing dropicles or particles to form a single layer when imaged.

### Production of hydrogel particles

Hydrogel precursor solution containing 10 wt% 8-arm PEG-vinylsulfone (JenKeM Technologies), 0.5 mg/ml biotin-PEG-thiol (Nanocs), 4 ng/ml maleimide labeled with Alexa Fluor 568 or 488 (Life Technologies), and 2 wt% dithiothreitol (Sigma-Aldrich) crosslinker was buffered in a 0.15M triethanolamine (Sigma-Aldrich) solution at pH = 5, and injected as the dispersed phase into the step emulsifier at 300 μL/hr, along with a continuous phase composed of Novec 7500 fluorinated oil and 1 wt% PicoSurf (Sphere Fluidics) at 1500 μL/hr to generate a water in oil emulsion.

At the device outlet, a second continuous phase containing 3 vol% triethylamine (Sigma-Aldrich) in Novec 7500 was introduced in a PTFE tubing (Zeus) via a Y junction connection (IDEX Health & Science) to increase the pH of the gel precursor droplets to pH 8 and initiate hydrogel crosslinking. Oil containing the organic base was injected using a Hamilton gastight syringe at 150 μL/hr. The residence time in the tubing between introduction of the organic base and the collection tube was around 4 min. All solutions were injected into the step emulsification device at defined flow rates using syringe pumps (Harvard Apparatus PHD 2000).

After incubating for 8 hr at room temperature, crosslinked particles were extracted from the emulsion by disrupting the emulsion with 20 wt% perfluoro-octanol (Sigma-Aldrich) in Novec 7500 oil at a 1:1 volume ratio. PBS was added immediately to swell and disperse the particles, followed by a few repeats of hexane (Sigma-Aldrich) extraction steps where the oil contents are extracted into a low-density hexane phase and pipetted out from the top. The particles were then filtered with Falcon 40 μm Cell Strainers (Corning) to remove oversized particle aggregates and debris and suspended in PBS supplemented with 0.1 wt% Pluronic F-127 (Sigma Aldrich). The density of the particle suspension was measured using standard cell counting techniques or a flow cytometer. The particles were stored at 4 °C for long term storage up to several months, and usually suspended at a density of 10^6^ particles /mL for ease of use in assays.

### Affinity binding on particles

Stored particle suspension was taken out of the fridge and homogenized by inversion. A calculated volume of suspension containing the desired number of particles was transferred to a protein LoBind microcentrifuge tube (Eppendorf) using Ultra Low Retention pipette tips (Corning) and spun down for 15 seconds using a benchtop centrifuge (Stellar Scientific). A translucent pellet of hydrogel particles could be seen at the bottom of the tube. The supernatant was discarded and replaced with 50 μl solutions of the target binding molecule at varying concentrations. The mixture was gently pipetted to dislodge and pellet and incubated on a rotating rack. After incubating for 1 hour, the particles were washed 4 times following similar centrifuge – solution exchange – mixing steps in PBS containing 0.1% Pluronic F-127.

### Dropicle formation

The particles were suspended in PBS supplemented with 0.1% Pluronic F-127 and pelleted for use. For enzymatic amplification assays, the fluorogenic substrate was also diluted in PBS with 0.1% Pluronic F-127 to a concentration of 100 μM, 5 μl of this substrate solution was gently mixed with the particles. An oil phase of Novec 7500 with 1% PicoSurf was immediately added, and this mixture was pipetted for 2 minutes at an approximate rate of 120 pipette aspiration-dispensing cycles per minute to create an emulsion of uniform dropicles.

The volume ratio of the aqueous phase and the oil phase is important for the emulsion quality. For all the enzymatic assays in this work, a pellet of 500,000 particles was mixed with 5 μL substrate solution and emulsified in 80 μL oil. 0.1% Pluronic F-127 in the aqueous phase was also important for the emulsion quality.

### Enzymatic affinity assays in particle templated droplets

#### β-gal amplification reaction

500,000 hydrogel particles were incubated with a 50 μl sample solution containing streptavidin-β-galactosidase (Sigma Aldrich) for 1 hour on a rotating rack set to 10 rotations per minute. Following incubation, particles were washed 4 times with PBS supplemented with 0.1% Pluronic F-127 and pelleted. 10 μL fluorescein-di-beta-galactopyranoside (FDG, Sigma Aldrich) solution at 100 μM in 0.1% Pluronic F-127 was mixed with the pellet. 80 μL of Novec 7500 with 0.5% PicoSurf was added to the pellet and immediately emulsified. The emulsion was incubated for 20 hours at room temperature covered in foil.

#### HRP amplification reaction

500,000 hydrogel particles were incubated with a sample solution containing Pierce High Sensitivity Streptavidin-HRP (Thermo Scientific) for 1 hour on a rotating rack set to 10 rotations per minute. Following incubation, particles were washed 4 times with PBS supplemented with 0.1% Pluronic F-127 and pelleted. Inside a dark room, 10 μL QuantaRed reaction solution (Thermo Scientific) mixed according to the manufacturer’s protocol was added the pellet. 80 μL of Novec 7500 with 0.5% PicoSurf was mixed with the pellet and immediately emulsified. The emulsion was immediately transferred to an imaging reservoir for signal readouts.

#### Affinity binding facilitated by active absorption

500,000 hydrogel particles were pelleted and washed in 500 μL ethanol for 3 times. After removing the ethanol supernatant after the third wash, the microcentrifuge tube containing the particles was placed inside a desiccator with the cap of the tube left open and kept in vacuum for 30 minutes in room temperature. Completely dried particles should be found at the bottom of the tube. Enzyme solutions were added directly to the dried particles, and immediately mixed with the rehydrating particles by gentle pipetting. The following incubation and emulsification steps are the same as above.

### Signal readout using fluorescence microscopy

An imaging reservoir was first filled with Novec 7500, followed by dropicles transferred by pipetting. The dropicles spread out into a single layer inside the reservoir since the height of the reservoir does not allow 2 particles stacked vertically. The reservoir containing the emulsion is then placed on the imaging stage of a Nikon Eclipse Ti2 Series equipped with a Photometrics Prime cMOS camera and scanned for 40 consecutive fields of views.

For β-gal/FDG/fluorescein reactions, each field of view was imaged in the FITC channel with 500 ms exposure to measure enzymatic signal in droplets, followed by 100ms in the TRITC channel to identify particles labelled with Alexa Fluor 568. For HRP/ADHP/resorufin reactions, each field of view was imaged in the TRITC channel with 40 ms exposure to measure enzymatic signal in droplets, followed by the FITC channel with 40 ms exposure to identify particles labelled with Alexa Fluor 488.

Image analysis algorithms were constructed in MATLAB to analyze the fluorescence signal of each dropicle by averaging signals over the particle. A positive signal is determined by thresholding 2.5x standard deviations above the mean of the background signals. Occasional occurrences of dropicles containing multiple hydrogel particles (<2%) were excluded from signal analysis.

To calculate the limit of detection (LOD), we measured the number of false positives in replicate (N=3) negative control samples. The detection threshold was set at MEAN+2.5SD (the sum of the mean fraction of false positives with 2.5 standard deviations of the fraction false positives), and the LOD was calculated by the intersect of the regression with the detection threshold.

The theoretical average enzyme per particle (AEP_theo_) was determined by Poisson distribution statistics (eq 1) where the average number of enzymes per particle λ = AEP_theo_ and the discrete occurrences of enzyme bound on particles X = x (x = 0, 1, 2, etc.) follows:

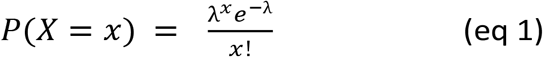

In the digital regime of the assay where the dropicles were either positive (x > 0) or negative (x = 0), the experimental average enzyme per particle (AEP_exp_) was calculated from the fraction of positive particles (f_p_) (eq 2)^53^:

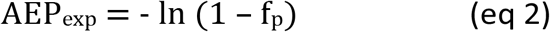

## Conflicts of interest

The authors have filed a patent application based in part on this work.

## Acknowledgements

This project is supported by the NSF-ERC PATHS-UP (award no. 1648451) and NSF-ERC TANMS (award no. 1160504) and UCLA SPORE in Prostate Cancer Support, Award Number P50 CA092131. We acknowledge the UCLA Nanofabrication Laboratory for access to microfabrication facilities and the UCLA Eli and Edythe Broad Center of Regenerative Medicine and Stem Cell Research.

## Notes

### Competing Interest Statement

The authors have declared no competing interest.

## References

1 A. S. Basu, SLAS Technol. Transl. Life Sci. Innov., 2017, 22, 369–386.

2 B. Rotman, Proc. Natl. Acad. Sci., 1961, 47, 1981–1991.

3 D. H. Wilson, D. M. Rissin, C. W. Kan, D. R. Fournier, T. Piech, T. G. Campbell, R. E. Meyer, M. W. Fishburn, C. Cabrera, P. P. Patel, E. Frew, Y. Chen, L. Chang, E. P. Ferrell, V. von Einem, W. McGuigan, M. Reinhardt, H. Sayer, C. Vielsack and D. C. Duffy, J. Lab. Autom., 2016, 21, 533–547.

4 T. Ono, T. Ichiki and H. Noji, Analyst, 2018, 143, 4923–4929.

5 Y. Rondelez, G. Tresset, K. V. Tabata, H. Arata, H. Fujita, S. Takeuchi and H. Noji, Nat. Biotechnol., 2005, 23, 361–365.

6 D. M. Rissin and D. R. Walt, J. Am. Chem. Soc., 2006, 128, 6286–6287.

7 D. M. Rissin, C. W. Kan, T. G. Campbell, S. C. Howes, D. R. Fournier, L. Song, T. Piech, P. P. Patel, L. Chang, A. J. Rivnak, E. P. Ferrell, J. D. Randall, G. K. Provuncher, D. R. Walt and D. C. Duffy, Nat. Biotechnol., 2010, 28, 595–599.

8 S. H. Kim, S. Iwai, S. Araki, S. Sakakihara, R. Iino and H. Noji, Lab Chip, 2012, 12, 4986–4991.

9 D. R. Walt, Lab Chip, 2014, 14, 3195–3200.

10 C. Wu, A. M. Maley and D. R. Walt, Crit. Rev. Clin. Lab. Sci., 2020, 57, 270–290.

11 S. S. Hwang, H. Chan, M. Sorci, J. Van Deventer, D. Wittrup, G. Belfort and D. Walt, Anal. Biochem., 2019, 566, 40–45.

12 A. Darlix, C. Hirtz, S. Thezenas, A. Maceski, A. Gabelle, E. Lopez-Crapez, H. De Forges, N. Firmin, S. Guiu, W. Jacot and S. Lehmann, BMC Cancer, 2019, 19, 110.

13 A. Bera, M. Subramanian, J. Karaian, M. Eklund, S. Radhakrishnan, N. Gana, S. Rothwell, H. Pollard, H. Hu, C. D. Shriver and M. Srivastava, PLoS One, 2020, 15, e0242141.

14 A. Vignoli, E. Muraro, G. Miolo, L. Tenori, P. Turano, E. Di Gregorio, A. Steffan, C. Luchinat and G. Corona, Cancers 2020, Vol. 12, Page 314, 2020, 12, 314.

15 P. Jarolim, P. P. Patel, M. J. Conrad, L. Chang, V. Melenovsky and D. H. Wilson, Clin. Chem., 2015, 61, 1283–1291.

16 J. Yang, K. Savvatis, J. S. Kang, P. Fan, H. Zhong, K. Schwartz, V. Barry, A. Mikels-Vigdal, S. Karpinski, D. Kornyeyev, J. Adamkewicz, X. Feng, Q. Zhou, C. Shang, P. Kumar, D. Phan, M. Kasner, B. López, J. Diez, K. C. Wright, R. L. Kovacs, P. S. Chen, T. Quertermous, V. Smith, L. Yao, C. Tschöpe and C. P. Chang, Nat. Commun., 2016, 7, 1–15.

17 A. J. Rivnak, D. M. Rissin, C. W. Kan, L. Song, M. W. Fishburn, T. Piech, T. G. Campbell, D. R. DuPont, M. Gardel, S. Sullivan, B. A. Pink, C. G. Cabrera, D. R. Fournier and D. C. Duffy, J. Immunol. Methods, 2015, 424, 20–27.

18 L. Song, D. W. Hanlon, L. Chang, G. K. Provuncher, C. W. Kan, T. G. Campbell, D. R. Fournier, E. P. Ferrell, A. J. Rivnak, B. A. Pink, K. A. Minnehan, P. P. Patel, D. H. Wilson, M. A. Till, W. A. Faubion and D. C. Duffy, J. Immunol. Methods, 2011, 372, 177–186.

19 R. Ahmad, L. Xie, M. Pyle, M. F. Suarez, T. Broger, D. Steinberg, S. M. Ame, M. G. Lucero, M. J. Szucs, M. MacMullan, F. S. Berven, A. Dutta, D. M. Sanvictores, V. L. Tallo, R. Bencher, D. P. Eisinger, U. Dhingra, S. Deb, S. M. Ali, S. Mehta, W. W. Fawzi, I. D. Riley, S. Sazawal, Z. Premji, R. Black, C. J. L. Murray, B. Rodriguez, S. A. Carr, D. R. Walt and M. A. Gillette, Sci. Transl. Med., 2019, 11, 8287.

20 F. Piraino, F. Volpetti, C. Watson and S. J. Maerkl, ACS Nano, 2016, 10, 1699–1710.

21 Y. Song, Y. Ye, S. H. Su, A. Stephens, T. Cai, M. T. Chung, M. K. Han, M. W. Newstead, L. Yessayan, D. Frame, H. D. Humes, B. H. Singer and K. Kurabayashi, Lab Chip, 2021, 21, 331–343.

22 J. U. Shim, R. T. Ranasinghe, C. A. Smith, S. M. Ibrahim, F. Hollfelder, W. T. S. Huck, D. Klenerman and C. Abell, ACS Nano, 2013, 7, 5955–5964.

23 V. Yelleswarapu, J. R. Buser, M. Haber, J. Baron, E. Inapuri and D. Issadore, Proc. Natl. Acad. Sci. U. S. A., 2019, 116, 4489–4495.

24 Y. Zhang and H. Noji, Anal. Chem., 2017, 89, 92–101.

25 C. Vielsack, R. E. Meyer, C. Cabrera, D. M. Rissin, T. G. Campbell, L. Chang, M. Reinhardt, D. R. Fournier, W. McGuigan, P. P. Patel, D. H. Wilson, V. von Einem, M. W. Fishburn, E. P. Ferrell, E. Frew, T. Piech, H. Sayer, D. C. Duffy, Y. Chen and C. W. Kan, J. Lab. Autom., 2015, 21, 533–547.

26 K. Akama, K. Shirai and S. Suzuki, Anal. Chem., 2016, 88, 7123–7129.

27 A. M. Maley, P. M. Garden and D. R. Walt, ACS Sensors, 2020, 5, 3037–3042.

28 K. Akama, K. Shirai and S. Suzuki, Electron. Commun. Japan, 2019, 102, 43–47.

29 T. Huynh, S. A. Byrnes, T. C. Chang, B. H. Weigl and K. P. Nichols, Analyst, 2019, 144, 7209–7219.

30 S. A. Byrnes, T. Huynh, T. C. Chang, C. E. Anderson, J. J. McDermott, C. I. Oncina, B. H. Weigl and K. P. Nichols, Anal. Chem., 2020, 92, 3535–3543.

31 R. Novak, Y. Zeng, J. Shuga, G. Venugopalan, D. A. Fletcher, M. T. Smith and R. A. Mathies, Angew. Chemie Int. Ed., 2011, 50, 390–395.

32 M. N. Hatori, S. C. Kim and A. R. Abate, Anal. Chem., 2018, 90, 9813–9820.

33 C. Y. Wu, M. Ouyang, B. Wang, J. de Rutte, A. Joo, M. Jacobs, K. Ha, A. L. Bertozzi and D. Di Carlo, Sci. Adv., 2020, 6, eabb9023.

34 G. Destgeer, M. Ouyang and D. Di Carlo, Anal. Chem., 2021, 93, 2317–2326.

35 J. M. de Rutte, J. Koh and D. Di Carlo, Adv. Funct. Mater., 2019, 29, 1900071.

36 Z. Huang, Kinetic Fluorescence Measurement of Fluorescein Di-j3-o-galactoside Hydrolysis by /3-Galactosidase: Intermediate Channeling in Stepwise Catalysis by a Free Single Enzyme1”, Matchett, 1991, vol. 30.

37 J. Hofmann and M. Sernetz, Anal. Biochem., 1983, 131, 180–186.

38 Z. Guan, Y. Zou, M. Zhang, J. Lv, H. Shen, P. Yang, H. Zhang, Z. Zhu and C. J. Yang, Biomicrofluidics, DOI:10.1063/1.4866766.

39 B. Porstmann, T. Porstmann, E. Nugel and U. Evers, J. Immunol. Methods, 1985, 79, 27–37.

40 B. Zhao, F. A. Summers and R. P. Mason, Free Radic. Biol. Med., 2012, 53, 1080–1087.

41 Y. Chen, A. Wijaya Gani and S. K. Y. Tang, Lab Chip, 2012, 12, 5093–5103.

42 P. Gruner, B. Riechers, L. A. Chacòn Orellana, Q. Brosseau, F. Maes, T. Beneyton, D. Pekin and J. C. Baret, Curr. Opin. Colloid Interface Sci., 2015, 20, 183–191.

43 G. Etienne, A. Vian, M. Biočanin, B. Deplancke and E. Amstad, Lab Chip, 2018, 18, 3903–3912.

44 P. Gruner, B. Riechers, B. Semin, J. Lim, A. Johnston, K. Short and J.-C. C. Baret, Nat. Commun., 2016, 7, 10392.

45 Y. Skhiri, P. Gruner, B. Semin, Q. Brosseau, D. Pekin, L. Mazutis, V. Goust, F. Kleinschmidt, A. El Harrak, J. B. Hutchison, E. Mayot, J. F. Bartolo, A. D. Griffiths, V. Taly and J. C. Baret, Soft Matter, 2012, 8, 10618–10627.

46 L. Chang, D. M. Rissin, D. R. Fournier, T. Piech, P. P. Patel, D. H. Wilson and D. C. Duffy, J. Immunol. Methods, 2012, 378, 102–115.

47 H. E. Muñoz, C. T. Riche, J. E. Kong, M. Van Zee, O. B. Garner, A. Ozcan and D. Di Carlo, ACS Sensors, 2020, 5, 385–394.

48 K. K. Brower, K. K. Brower, C. Carswell-Crumpton, S. Klemm, B. Cruz, G. Kim, S. G. K. Calhoun, L. Nichols, P. M. Fordyce, P. M. Fordyce, P. M. Fordyce and P. M. Fordyce, Lab Chip, 2020, 20, 2062–2074.

49 K. K. Brower, M. Khariton, P. H. Suzuki, C. Still, G. Kim, S. G. K. Calhoun, L. S. Qi, B. Wang and P. M. Fordyce, Anal. Chem., 2020, 92, 13262–13270.

50 K. Takeda, Y. Fujimura and F. Jimma, Cytom. Res., 2011, 20, 43–50.

51 G. Destgeer, M. Ouyang, C. Y. Wu and D. Di Carlo, Lab Chip, 2020, 20, 3503–3514.

52 M. Li, M. van Zee, C. T. Riche, B. Tofig, S. D. Gallaher, S. S. Merchant, R. Damoiseaux, K. Goda and D. Di Carlo, Small, 2018, 14, 1803315.

53 D. M. Rissin, D. R. Fournier, T. Piech, C. W. Kan, T. G. Campbell, L. Song, L. Chang, A. J. Rivnak, P. P. Patel, G. K. Provuncher, E. P. Ferrell, S. C. Howes, B. A. Pink, K. A. Minnehan, D. H. Wilson and D. C. Duffy, Anal. Chem., 2011, 83, 2279–2285.

