## Supplementary material for "Counting of Enzymatically Amplified Affinity Reactions in Hydrogel Particle-Templated Drops": Fig. S

Please do not adjust margins



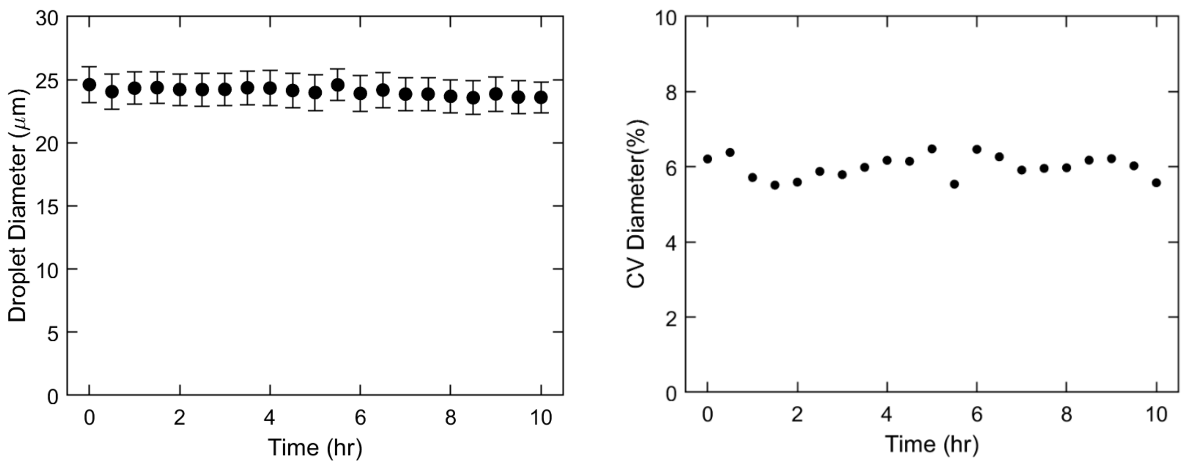


**Figure. S1** The mean diameter of droplets produced over a 10-hour period using the step emulsifier microfluidic device remains consistent, with CV values around 6%. (n = 150-300 for each 30-minute time point).

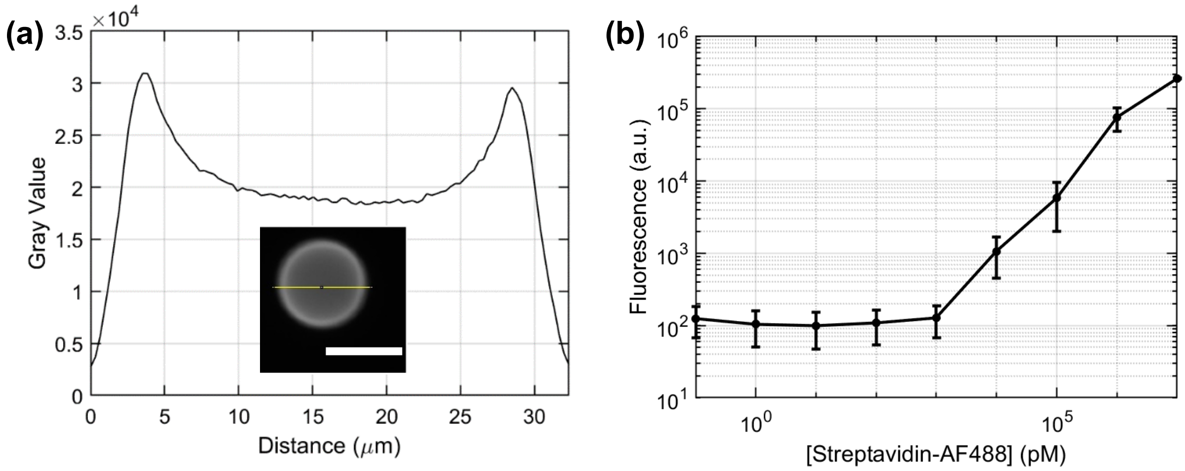


**Figure. S2** Streptavidin binding to biotinylated particles. **(a)** Fluorescence signal is localized to the outer surface of the hydrogel particle, forming a bright edge on the boundary of the particle in a 1D fluorescence intensity slice. **(b)** The integrated fluorescence signal from particles with bound streptavidin-Alexa Flour 488 after incubation are linearly correlated to the concentration of the streptavidin solution across 5 orders of magnitudes. The lowest resolvable signal was around 1 nM, representing the limit of detection for an unamplified affinity assay on the particles using our microscopy setup. (n = 10,000 for each streptavidin concentration.)



Please do not adjust margins

Please do not adjust margins



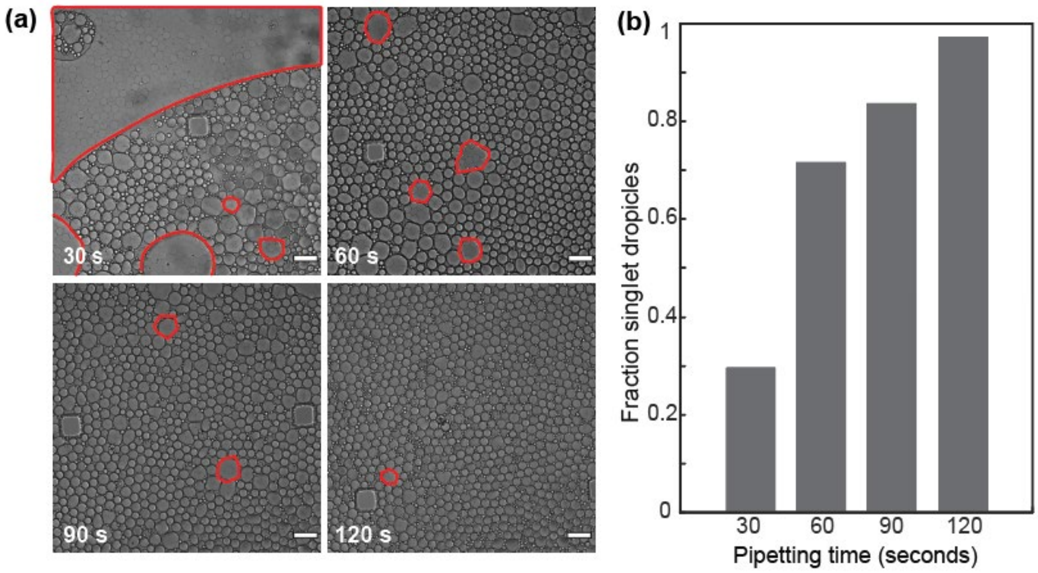


**Figure. S3** Optimizing the quality of dropicle formation with increased pipetting time. **(a)** Bright field images showing the formation of dropicles at 30 second intervals of vigorous pipetting. The number of droplets containing multiple hydrogel particles (highlight by red contour) decreased with increased pipetting time. Scale bar = 100 µm. **(b)** The fraction of singlet dropicles (droplets templated by only one particle) increased with time, approaching 100% after 120 seconds of pipetting.

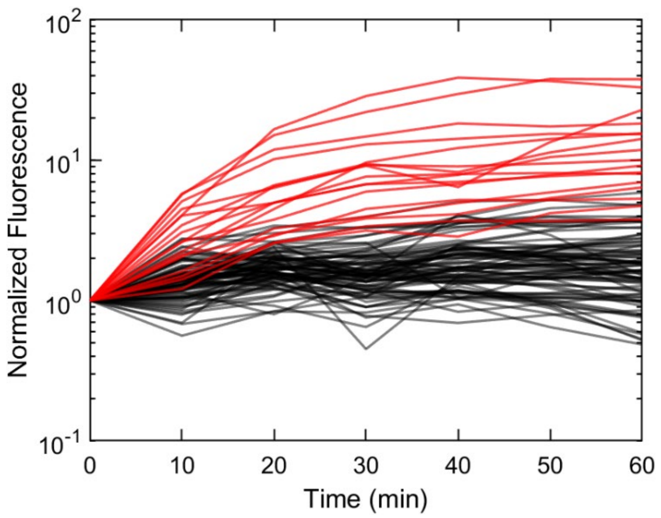


**Figure. S4** Signals of dropicles loaded with the HRP/ADHP/resorufin system analyzed at 10-minute intervals indicating the enzymatic amplification completed at around 30 minutes. The fluorescence signals of each dropicle was normalized against their starting fluorescence at t=0. Red lines refer to signals from positive dropicles (containing particles bound with at least 1 HRP enzyme), black lines correspond to negative dropicles. (n = 101)


This journal is © The Royal Society of Chemistry 20xx

Please do not adjust margins

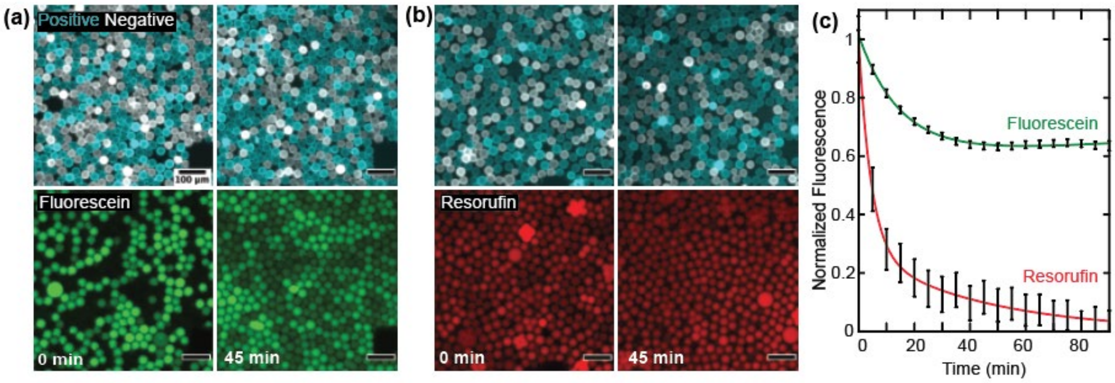


**Figure. S5** Transport of resorufin and fluorescein in dropicles. **(a)** Fluorescence signals from fluorescein containing dropicles, the location of which indicated by the cyan color filter on the top images, transported towards negative dropicles during a 45-minute incubation. Scale bar = 100 μm. **(b)** Fluorescence signals from resorufin containing dropicles transported towards negative dropicles during a 45-minute incubation. **(c)** Normalized fluorescence intensities over 90 minutes observation showing the transport of fluorescein is slower than resorufin, and ~65% of fluorescein signal was retained at equilibrium. Error bars indicate the standard deviation of all dropicles observed.

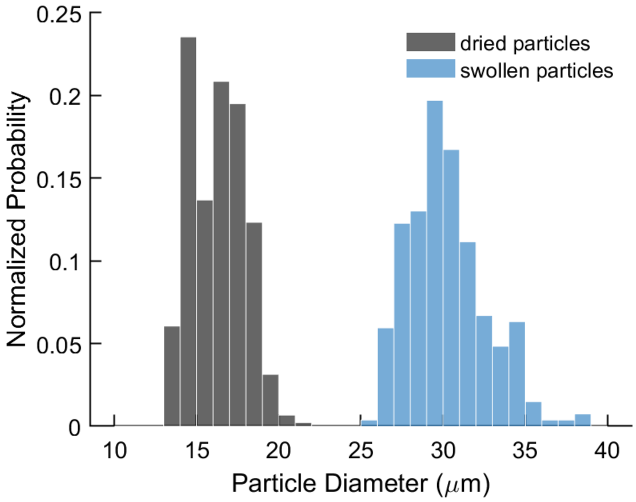


**Figure. S6** Change of particle size during the rehydration process of active absorption. Dried particles ($d_1$ = 16.3 ± 1.6 μm, $n_1$ = 446) swelled into hydrated particles ($d_2$ = 30.3 ± 2.4 μm, $n_2$ = 269), yielding a 6.4-fold in the total volume of the spherical hydrogel particles.
